# The ubiquitin E3 ligase Huwe1 facilitates viral and self RNA sensing by RIG-I-like receptors

**DOI:** 10.1101/2025.03.27.645708

**Authors:** Timo Oosenbrug, Dennis Gravekamp, Adán Pinto-Fernández, Stefania Crotta, Katja Finsterbusch, Sander B. van der Kooij, Laurens R. ter Haar, Yanniek Asschert, Gilaine F.E. Kleian, Benedikt M. Kessler, Andreas Wack, Annemarthe G. van der Veen

## Abstract

RIG-I-like receptors (RLRs) are cytoplasmic RNA sensors that promote type I and type III interferon (IFN) production in response to RNA ligands of viral or endogenous origin. The RLR pathway is tightly regulated by dynamic post-translational modifications, including ubiquitination. Huwe1 is a HECT domain-containing giant ubiquitin E3 ligase that has not been implicated in the RLR or IFN pathway. Here, we investigated whether Huwe1 is required for type I IFN induction downstream of RLRs. We demonstrate that loss of Huwe1 severely attenuates the expression of IFN-β, IFN-λ1 and IFN-stimulated genes (ISGs) in ADAR1-deficient human cells and primary murine bone-marrow derived macrophages, in which unedited self RNAs that serve as RLR ligands accumulate. In addition, depletion of Huwe1 reduces the induction of type I and III IFNs upon transfection with synthetic viral RNA mimetics or infection with a picornavirus. Using proteomics, we identified several putative Huwe1 substrates, which include key components of the RLR pathway (MAVS, TRAFs). We demonstrate that these substrates interact with Huwe1 and that Huwe1 is essential for the activity of TRAF5 in type I IFN induction. Collectively, our results put Huwe1 on the map as an important ubiquitin E3 ligase in the RLR pathway and provide new insights into ubiquitin-dependent regulation of cell-intrinsic antiviral immune pathways.

## Introduction

Nucleic acid sensing and the type I interferon (IFN) response mediate antiviral immunity against viruses. Cytosolic and endosomal DNA and RNA sensors detect virus-derived nucleic acids and initiate a downstream signalling cascade that leads to the production and secretion of type I and type III IFNs^1–3^. These antiviral cytokines activate their respective receptors in an autocrine and paracrine manner, which induces signalling via Janus kinase (JAK) and signal transducer and activator of transcription (STAT1/2), leading to the upregulation of hundreds of IFN-stimulated genes (ISGs) that mediate antiviral defence^4, 5^. While all nucleated cells can produce and respond to type I IFNs, the expression of the type III IFN receptor is restricted to epithelial cells, hence type III IFNs protect epithelial barrier surfaces against viral infection^6^.

The family of RIG-I-like receptors (RLRs) includes three related RNA helicases that detect distinct viral (non-self) RNA ligands in the cytosol of infected cells, namely retinoic acid-inducible gene I (RIG-I), melanoma differentiation-associated protein 5 (MDA5), and laboratory of genetics and physiology 2 (LGP2), the least well-understood member of the RLR family^7–9^. RIG-I and MDA5 both contain two N-terminal CARD domains that are important for oligomerization and signalling to the downstream adaptor mitochondrial antiviral signalling (MAVS)^10^. Activation and oligomerization of MAVS subsequently leads to the recruitment and activation of TNF receptor-associated factor (TRAF) ubiquitin E3 ligases, Tank-binding kinase 1 (TBK1) and the IκB kinase (IKK) complex. This, in turn, promotes nuclear translocation of the transcription factors interferon regulatory factor 3 (IRF3) and NF-κB, which initiates the transcription of type I and type III IFNs and pro-inflammatory cytokines, respectively^11^. LGP2 lacks CARD domains and cannot signal via MAVS. Instead, LGP2 facilitates MDA5 activation by nucleating MDA5 filament formation on double-stranded RNA (dsRNA)^12–14^, by optimising the length of the MDA5 filaments, and/or by serving as a ‘road block’ that prevents MDA5 from running of the filament^14–17^. Conversely, LGP2 restricts RIG-I activation, although the precise mechanism underlying this function remains poorly defined^18–21^.

While type I and type III IFNs are essential to establish a successful antiviral immune response, chronic, and/or excessive production of IFNs is detrimental and causes immunopathology. This is best exemplified by the type I interferonopathies, a spectrum of inherited autoinflammatory disorders characterized by chronic type I IFN production in the absence of a viral agent, due to genetic defects^22^. Amongst such genetic defects are mutations in the RNA editing enzyme adenosine deaminase acting on RNA 1 (ADAR1), which changes adenosines into inosines, in particular in repetitive regions with high intramolecular complementarity. Loss of A-to-I editing leads to the accumulation of unedited, endogenous RNA molecules with a high degree of base-pairing that resemble viral dsRNA^23–27^. Such immunostimulatory self RNA molecules are detected by the RNA sensors MDA5 and LGP2^28–32^. The presence of LGP2 is crucial to mount an IFN response to unedited, self RNA following loss of ADAR1 activity. Genetic ablation of LGP2 in human cell lines or mouse models prevents type I IFN induction in ADAR1-deficient context^31, 32^.

Type I IFN production and signalling requires a fine-tuned balance to ensure an effective antiviral response while avoiding autoinflammation and immunopathology. Regulatory mechanisms that keep nucleic acid sensing and IFN signalling in check include post-translational modifications, such as ubiquitination via the E1-E2-E3 enzymatic cascade, where E3 enzymes determine substrate specificity^33, 34^. Ubiquitination positively or negatively regulates various steps in the RLR pathway^34–36^. Several components of the RLR pathway, including the RLRs themselves and MAVS, are ubiquitinated to promote and stabilize their active conformation^11, 37–40^. TRAF E3 ligases, including TRAF2, -3, -5, and -6, form ubiquitin chains to attract and coordinate the activity of downstream kinases IKK and TBK1, although there is a certain degree of controversy in the field as to which of these TRAF proteins are critical in RLR signaling due to redundancy and cell type specificity^41–46^. Conversely, ubiquitination events that promote proteasomal degradation of players in the RLR pathway facilitate the resolution of signalling^34, 47^. Given the importance of keeping type I IFN induction at bay, additional mechanisms that regulate RLR activation likely exist.

HECT, UBA, and WWE domain-containing E3 ubiquitin protein ligase 1 (Huwe1, also known as HECTH9, Mule, ARF-BP1, Lasu1, Ureb1) is an evolutionary conserved, 482 kDa ubiquitin E3 ligase that mediates K6-, K11-, K48-, and to a lesser extent K63-linked ubiquitination^48, 49^. Huwe1 is involved in cell proliferation, DNA damage, cancer, and cellular stress, by ubiquitinating proteins such as Mcl-1, Chk1, Myc and p53^50–53^. In addition, mutations in Huwe1 have been associated with intellectual neurodevelopmental disorders^54–56^. Knowledge on its role in innate immune signalling is limited. Huwe1 regulates IL1β-induced NF-κB activation by building K48-linked ubiquitin branches onto K63-chains formed by TRAF6, thereby protecting TRAF6 with such K48-K63-branched linkages against deubiquitination and allowing sustained signalling to occur, ultimately promoting NF-κB activity^57^. In addition, Huwe1 has been implicated in inflammasome activation and defence against bacterial infection in mice^58^. Whether Huwe1 plays a role in nucleic acid sensing or the type I IFN response, is unknown.

Here, we find that Huwe1 is broadly required for RLR activation by self RNA ligands in multiple human cell lines and in primary bone marrow-derived macrophages. In addition, Huwe1 facilitates type I IFN induction in response to viral RNA mimetics or RNA virus infection in human cells. Using a mass spectrometry-based approach to identify Huwe1 substrates and biochemical experiments, we uncover that Huwe1 interacts with multiple potential substrates within the RLR pathway, including MAVS and TRAF proteins. We discovered that Huwe1 is important for the activity of TRAF5 in type I IFN induction. Finally, we find that co-depletion of Huwe1 and UBR5, another HECT E3 ligase that shares substrates and cooperates with Huwe1, further abrogates TRAF5 activity. Altogether, Huwe1 controls signalling events downstream of RLRs and regulates RLR activation in response to RNA ligands of viral and endogenous origin, providing new avenues for treatment of autoinflammatory and infectious diseases.

## Results

### Huwe1 is required to elicit a type I IFN response following self RNA sensing by RIG-I like receptors in ADAR1 deficiency

To determine whether the ubiquitin E3 ligase Huwe1 contributes to RLR activation and type I IFN induction, we first focused on a model system of self RNA sensing and sterile inflammation. Loss of ADAR1 expression, most notably its p150 isoform, leads to the accumulation of unedited, endogenous dsRNA that activates MDA5 and LGP2 and induces a sterile type I IFN response^31, 32^. Depletion of ADAR1 by siRNAs from LGP2-expressing human embryonic kidney 293 cells (HEK293-L, see methods and ^31^) resulted in robust upregulation of transcripts encoding IFN-β (type I IFN), IFN-λ1 (type III IFN) and IFIT1 (a prototype ISG), as determined by RT-qPCR (Fig. 1A). As expected, co-depletion of MDA5 completely blocked the upregulation of these transcripts. Importantly, depletion of Huwe1 using three individual siRNAs markedly reduced the upregulation of transcripts encoding IFN-β, IFN-λ1 and IFIT1 in ADAR1-deficient cells (Fig. 1A). Consistently, depletion of Huwe1 lowered the amount of bioactive type I IFN that was secreted (Fig. 1B). Moreover, reduced Huwe1 expression inhibited phosphorylation of STAT1, a key transcription factor that acts downstream of the type I and type III IFN receptors, and prevented upregulation of ISG60 protein expression, another prototype ISG (Fig. 1C). Knockdown efficiency of siRNA targets was validated by RT-qPCR and immunoblotting (Fig. S1A; Fig. 1C). Using an independent pool of Huwe1-targeting siRNAs, we also observed a reduced type I IFN response upon loss of ADAR1 (Fig. S1B-C). Finally, inhibition of Huwe1 activity using two previously described small molecule inhibitors (BI8622 and BI8626)^59^ led to similar outcomes (Fig. S1D). Taken together, these findings demonstrate that Huwe1 promotes type I IFN induction and signalling following loss of ADAR1.

**Figure 1:**
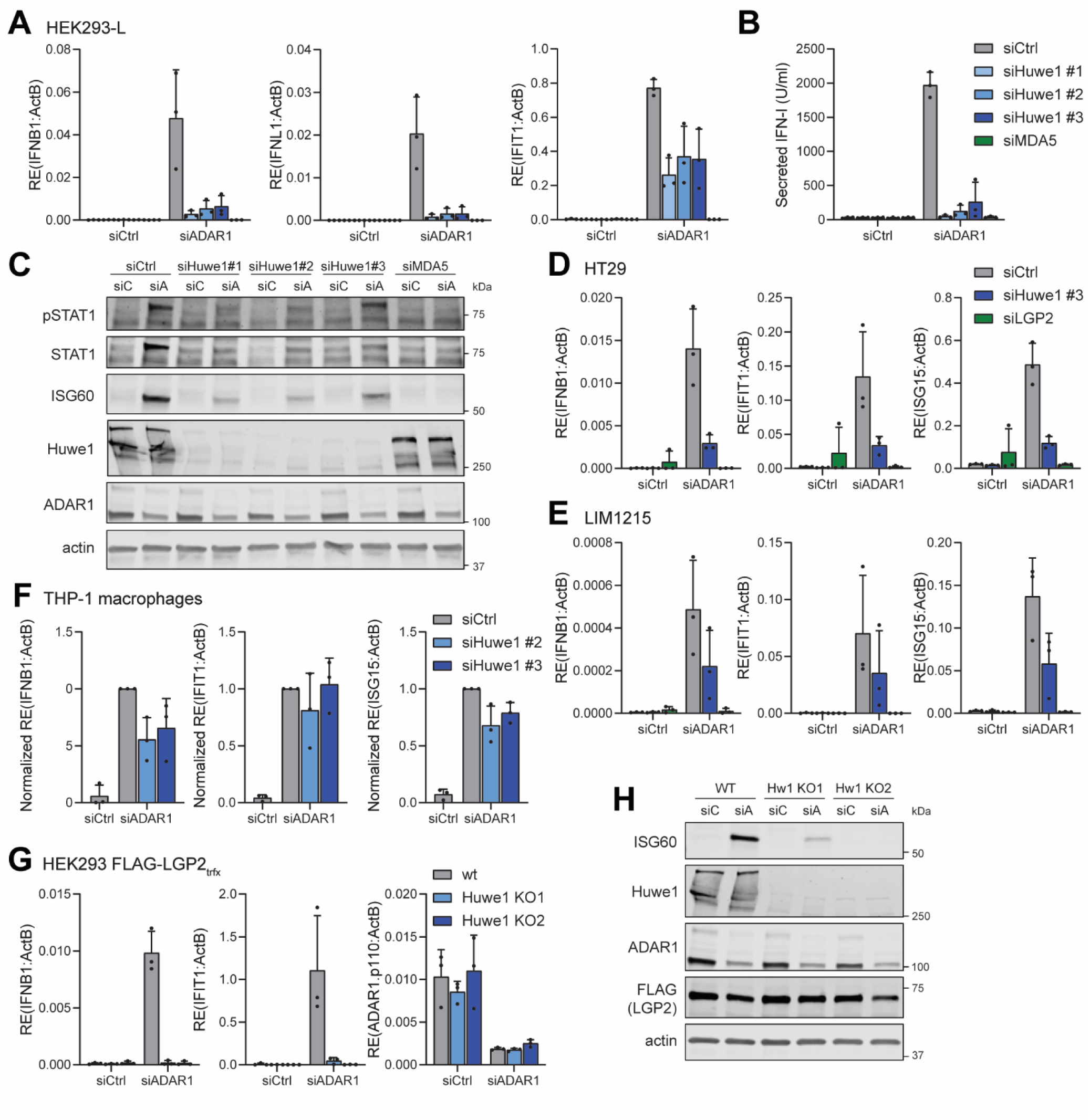
Huwe1 is required for type I and type III IFN induction in ADAR1-deficient cells. **A-C)** HEK293 cells that stably express FLAG-LGP2 (HEK293-L) were transfected with the indicated siRNAs. Cells and cell culture supernatants were harvested 72 h post-transfection. **A)** RT-qPCR analysis was used to monitor the type I and type III IFN response (IFN-β, IFN-λ1, and IFIT1 transcripts). Transcripts were normalised to ACTB. **B)** A bioassay was performed on collected cell culture supernatants to measure secreted IFN-I levels. **C)** Protein lysates were subjected to SDS-PAGE and immunoblotting using the indicated antibodies. siC, siCtrl; siA, siADAR1. **D-E)** The colorectal cancer cell lines HT29 (E) and LIM1215 (F) were transfected with the indicated siRNAs. Cells were harvested 72 h post-transfection and RT-qPCR analysis was used to monitor the induced type I IFN response. Transcripts were normalised to ACTB. **F)** PMA-differentiated THP-1 macrophages were transfected with the indicated siRNAs. Cells were harvested 96 h post-transfection and RT-qPCR analysis was used to monitor the induced type I IFN response. Transcripts were normalised to ACTB and are displayed relative to siCtrl/siADAR1. **G-H)** Wild type (WT) and Huwe1 knockout (KO) HEK293 cells were transfected with the indicated siRNAs and 8 h later with a vector encoding FLAG-LGP2. Cells were harvested 72 h post siRNA-transfection. **G)** RT-qPCR analysis was used to monitor the type I IFN response (IFN-β and IFIT1) and ADAR1 knockdown efficiency. Transcripts were normalised to ACTB. **H)** Protein lysates were subjected to SDS-PAGE and immunoblotting using the indicated antibodies. siC, siCtrl; siA, siADAR1. Data are presented as means ± s.d. from n=3 experiments (A, B, D, E-G) or are representative data (n=3 for C; n=2 for H).

We examined the impact of Huwe1 depletion in several other human cell lines. Huwe1 depletion also blocked MDA5/LGP2-dependent type I IFN induction upon loss of ADAR1 in the human colorectal cancer cell lines HT29 and LIM1215 (Fig. 1D-E; Fig. S1E-F). In addition, siRNA-mediated knockdown of Huwe1 caused a modest reduction in IFN-β and ISG transcripts in ADAR1-depleted THP-1 macrophages, despite limited knockdown efficiency in these cells (Fig. 1F; Fig. S1G). Thus, Huwe1 is required for type I IFN induction following RLR activation by unedited self RNA across multiple cell types.

To further substantiate above observations, we knocked out Huwe1 in HEK293 cells using CRISPR/Cas9-mediated genome engineering, to eliminate any residual enzymatic activity of Huwe1 due to incomplete siRNA-mediated knockdown. We obtained two Huwe1 knockout (KO) clones using distinct gRNAs. PCR analysis using primers that flank each gRNA binding site revealed targeted indel generation (Fig. S1H) and immunoblotting confirmed the concomitant loss of Huwe1 protein expression (Fig. S1I). When we depleted ADAR1 in wild type and Huwe1 KO cells, we observed reduced expression of IFN-β and IFIT1 transcripts as well as ISG60 protein in both Huwe1 KO clones (Fig. 1G and 1H). We conclude that Huwe1 is required for type I IFN induction following activation of LGP2/MDA5 by immunostimulatory self RNA in ADAR1-deficient cells.

### Huwe1 impacts on RLR signalling downstream of MDA5 oligomerization and upstream of IRF3 nuclear translocation

To dissect at which step of the RLR pathway Huwe1 exerts its role, we first tested whether Huwe1 is required for receptor activation. Activation of MDA5 involves its oligomerization along dsRNA ligands^60^, which in context of ADAR1 deficiency critically relies on expression of LGP2. These MDA5 aggregates can be detected by semi-denaturing detergent agarose-gel electrophoresis (SDD-AGE)^37, 61^. As expected, siRNA-mediated knockdown of ADAR1 in HEK293-L cells led to the formation of endogenous MDA5 aggregates (Fig. 2A). Co-depletion of Huwe1 did not inhibit MDA5 aggregation and perhaps even slightly increased aggregation. We confirmed these findings using ADAR1 KO HEK293 cells that are engineered to overexpress LGP2 in a doxycycline-inducible manner^31^. In line with previous findings, the addition of doxycycline to these cells induced LGP2 expression, which promoted MDA5 oligomerization as well as type I and III IFN expression (Fig. S2A-B). MDA5 oligomerization was unaffected by Huwe1 depletion in these cells (Fig. S2A), whereas the loss of Huwe1 did inhibit IFN-β and IFN-λ1 expression (Fig. S2A-B). Collectively, these findings indicate that Huwe1 is not required to promote the activation-induced oligomerization of MDA5 and instead acts downstream of receptor activation.

**Figure 2:**
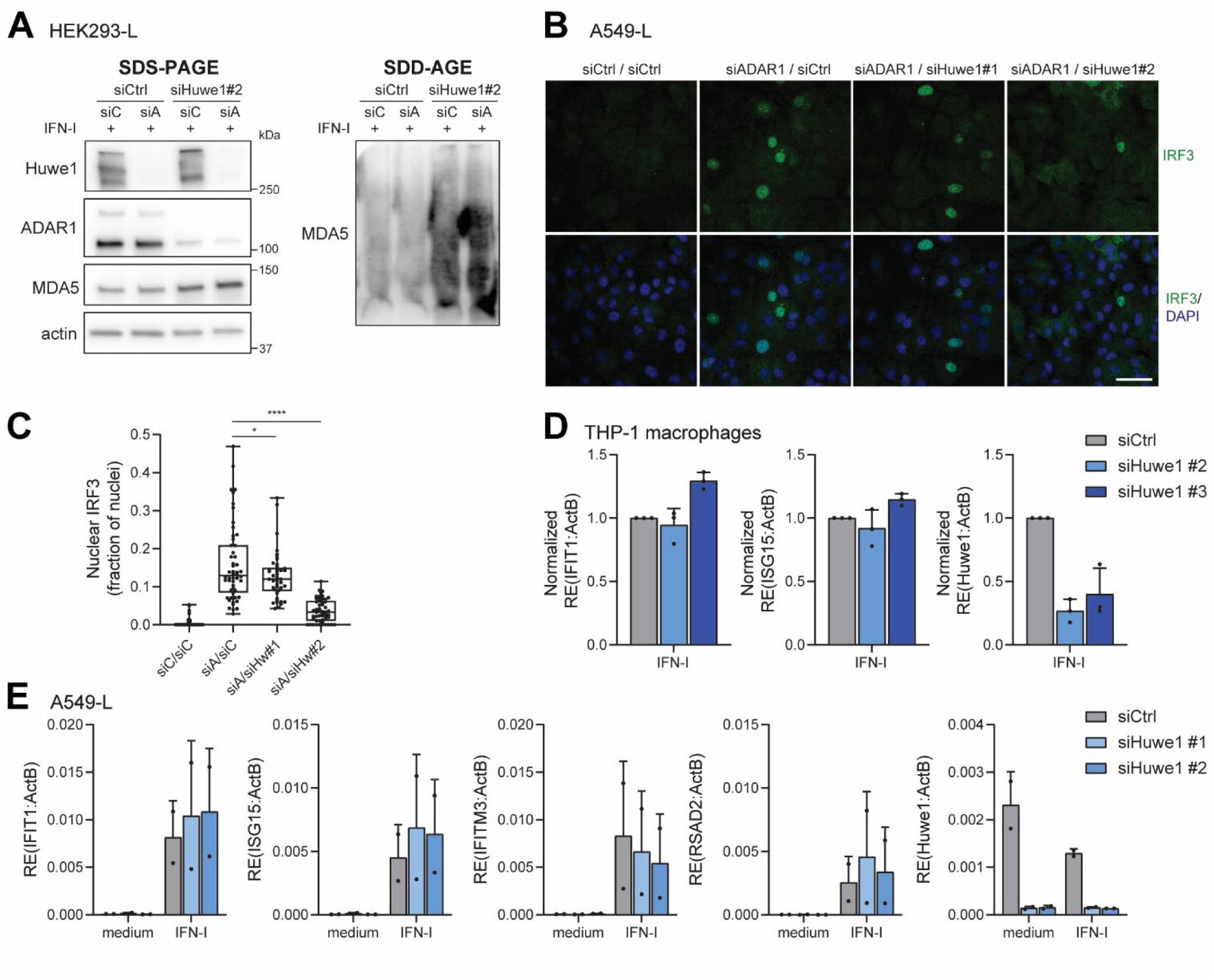
Huwe1 impacts on the RLR pathway downstream of receptor oligomerization and upstream of the nuclear translocation of IRF3. **A)** HEK293 cells that stably express FLAG-LGP2 (HEK293-L) were transfected with the indicated siRNAs for 72 h. During the last 24 h, recombinant type I IFN (1000 U/ml; IFN-I) was added to upregulate endogenous MDA5 protein expression. Protein lysates were subjected to SDD-AGE and SDS-PAGE, followed by immunoblotting using the indicated antibodies, to determine MDA5 oligomerization and total protein levels, respectively. siC, siCtrl; siA, siADAR1. **B)** The lung epithelial cell line A549, engineered to stably express FLAG-LGP2 (A549-L), was transfected with the indicated siRNAs for 72 h. Cells were stained with anti-IRF3 (green) and nuclei were stained with DAPI (blue). Scale bar indicates 50 µm. **C)** Quantification of immunofluorescence data presented in (B). Total and IRF3-positive nuclei were counted using CellProfiler and plotted as fraction of IRF3-positive nuclei per field of view. The boxplot indicates the interquartile range as a box, the median as a central line, and the whiskers extend from the minimum to the maximum value (n=2, >40 fields of view per experimental condition). Statistical analysis was performed using a one-way ANOVA test with a Dunnett’s post hoc test for multiple comparisons. *, p<0.05; ****, p<0.0001; siC, siCtrl; siA, siADAR1; siHw, siHuwe1. **D-E)** PMA-differentiated THP-1 macrophages (D) and A549-L cells (E) were treated for 16h with recombinant type I IFN (1000 U/ml; IFN-I). RT-qPCR was performed to monitor ISG transcription and Huwe1 knockdown efficiency. Transcripts were normalised to ACTB. In (D), data is displayed relative to siCtrl. Data are presented as means ± s.d. from n=3 (D) or n=2 (E) experiments or are representative data (n=3 for A; n=2 for B).

Activated RLRs trigger signalling that culminates in the activation and nuclear translocation of the transcription factor IRF3^62^. We used immunofluorescence to test whether Huwe1 impacts on this translocation event. Depletion of ADAR1 in an LGP2-expressing lung epithelial cell line (A549-L), which is more suitable for microscopy-based experiments than HEK293, clearly led to accumulation of IRF3 in the nucleus (Fig. 2B). Co-depletion of Huwe1 reduced nuclear translocation of IRF3, as quantified by the fraction of IRF3-positive nuclei (Fig. 2B-C). This suggests that Huwe1 impacts on IFN induction downstream of RLR activation but prior to IRF3 translocation. In line with this, upregulation of ISG transcripts following activation of the type I IFN receptor (IFNAR) by recombinant type I IFN treatment was unaffected by Huwe1 depletion in multiple cell types (Fig. 2D-E). Altogether, these findings indicate that Huwe1 enhances signalling at a step downstream of RLR receptor activation to promote proper IRF3 activation.

### Huwe1 promotes type I IFN induction upon RNA virus infection

We next investigated whether Huwe1, by modulating signalling downstream of receptor activation, also affects the type I IFN response induced by viral RNA. To this end, wild type and Huwe1 KO HEK293 cells were transfected with two synthetic ligands that mimic virus-derived dsRNAs, namely high molecular weight (HMW) poly(I:C) and small triphosphorylated hairpin RNA (3p-hpRNA) that activate MDA5 and RIG-I, respectively. RLR activation by either ligand induced expression of IFN-β-, IFIT1-, and ISG15-encoding transcripts, which was reduced in Huwe1 KO cells (Fig. 3A). In addition, in cells transfected with a reporter construct that encodes firefly luciferase under control of the IFN-β promoter, both HMW poly(I:C) and low molecular weight (LMW) poly(I:C) (which preferentially activates RIG-I), triggered less IFN-β promoter activity in Huwe1 KO clones compared to wild type cells (Fig. S3A). Huwe1 depletion by siRNAs in THP-1 macrophages also inhibited IFN-β mRNA upregulation by HMW poly(I:C) or 3p-hpRNA, albeit not to an extend that negatively affects subsequent ISG mRNA expression, which may be explained by incomplete Huwe1 knockdown (Fig. 3B). To determine whether Huwe1 also impacts on IFN induction following cGAS/STING activation, we treated THP-1 macrophages with two different dsDNA ligands. This led to a slight reduction in IFN-β induction in Huwe1-depleted cells (Fig. S3B), which may either reflect a different role for Huwe1 in the cytosolic DNA sensing pathway or Huwe1-dependent regulation of a shared signalling intermediate. Finally, we infected A549-L cells that were transfected with control or Huwe1-targeting siRNAs with Mengovirus. The viral RNAs produced by this cardiovirus from the *Picornaviridae* family are sensed by LGP2 and MDA5^63^. For optimal type I IFN induction, we made use of an attenuated virus with a mutation in the L protein (L^Zn^), which prevents the virus from antagonizing type I IFN expression^64^. Mengovirus L^Zn^ infection induced robust levels of IFN-β as well as IFN-λ1 expression in control cells, which was reduced in Huwe1-depleted cells (Fig. 3C; S3C). Transcription of the Mengovirus L region was unaffected (Fig. 3C), indicating that Huwe1 does not directly impact on viral replication and that the reduction in type I and III IFN expression was not due to reduced exposure to MDA5/LGP2 ligands. We conclude that, in addition to promoting the response to immunostimulatory self RNA, Huwe1 also facilitates type I IFN expression when RLRs are activated by diverse viral RNA ligands.

**Figure 3:**
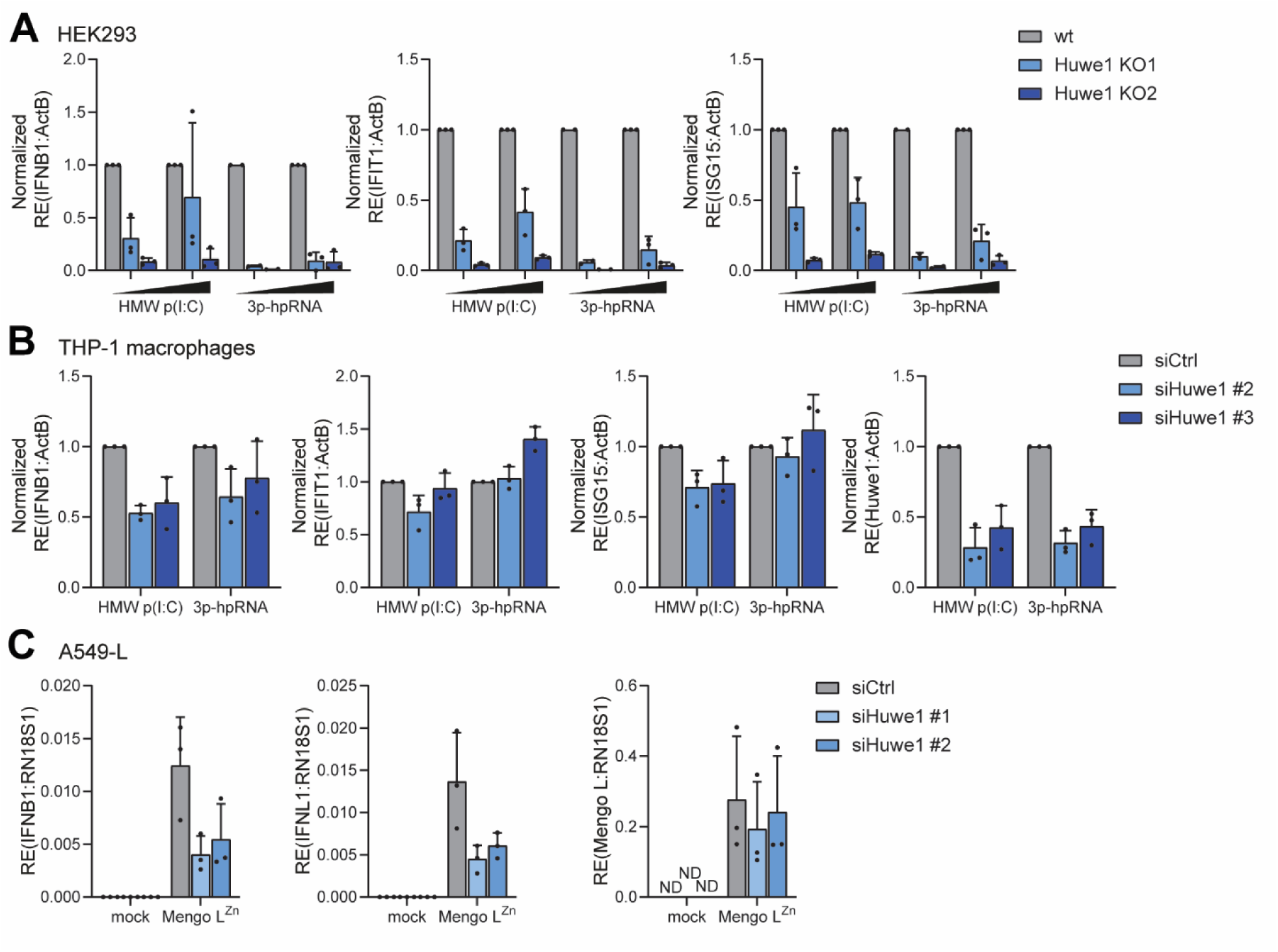
Loss of Huwe1 diminishes type I and type III IFN induction upon RLR activation by viral RNA mimetics or viral infection. **A)** Wild type (WT) and Huwe1 knockout (KO) HEK293 cells were transfected for 16 h with 100 or 200 ng of high molecular weight poly(I:C) (HMW p(I:C)) or 50 or 100 ng of a small triphosphorylated hairpin RNA (3p-hpRNA). RT-qPCR analysis was used to monitor the type I IFN response. Transcripts were normalised to ACTB and are displayed relative to WT cells. **B)** PMA-differentiated THP-1 macrophages were transfected with the indicated siRNAs. After 48 h, cells were transfected for an additional 16 h with 0.5 µg of HMW p(I:C) or 0.25 µg of 3p-hpRNA. RT-qPCR analysis was used to monitor the type I IFN response and Huwe1 knockdown efficiency. Transcripts were normalised to ACTB and are displayed relative to siCtrl. **C)** A549 cells that stably express FLAG-LGP2 (A549-L) were transfected with the indicated siRNAs. After 48 h, cells were infected with Mengovirus with a mutation that reduces viral antagonism of the type I IFN response in the L protein (L^Zn^) at MOI = 0.25 for 16 h. RT-qPCR analysis was used to monitor the type I IFN response and upregulation of Mengovirus RNA. Transcripts were normalised to RN18S1. ND, not detected. Data are presented as means ± s.d. from n=3 experiments (A-C).

### Loss of Huwe1 abrogates type I IFN induction upon depletion of ADAR1 in primary murine bone-marrow derived macrophages

To consolidate our findings in primary murine cells, we obtained bone marrow from Huwe1^fl/Y^R26-CreERT2 mice (Cre^+^), in which Huwe1 can be functionally depleted upon the tamoxifen (4-OHT)-induced expression of Cre recombinase, which excises part of the Huwe1 gene that encodes the catalytic C-terminal HECT domain^65^. Bone marrow from CreERT2 negative (Cre^-^) Huwe1^fl/Y^ mice served as control. Tamoxifen treatment during the M-CSF-mediated differentiation of bone-marrow derived macrophages (BMDMs) was well-tolerated and did not have an adverse effect on the expression of differentiation markers (Fig. S4A). After differentiation, BMDMs were treated with cell-penetrating phosphoroamidate morpholino oligomers (Vivo-PMOs) to intervene with ADAR1 expression by inducing exon skipping (Fig. 4A). Loss of ADAR1 led to the induction of a type I IFN response in control BMDMs, which was selectively lost in tamoxifen-treated Cre^+^ cells (i.e. Huwe1-deficient) (Fig. 4B-C). RT–qPCR using primers specific for the floxed region confirmed efficient Huwe1 recombination in the tamoxifen-treated Cre^+^ cells (Fig. 4B). Loss of full-length Huwe1 protein, but not a lower molecular weight product, was confirmed by immunoblotting (Fig. 4C). In contrast to our observations in human cell lines, the induction of a type I IFN response following transfection of HMW poly(I:C) or 3p-hpRNA to stimulate MDA5 or RIG-I, respectively, was not affected by the loss of Huwe1 in BMDMs (Fig. S4B), possibly due to cell type or species-specific differences. We conclude that in primary murine BMDMs Huwe1 is required for sensing of unedited immunostimulatory RNA in ADAR1-deficient cells.

**Figure 4:**
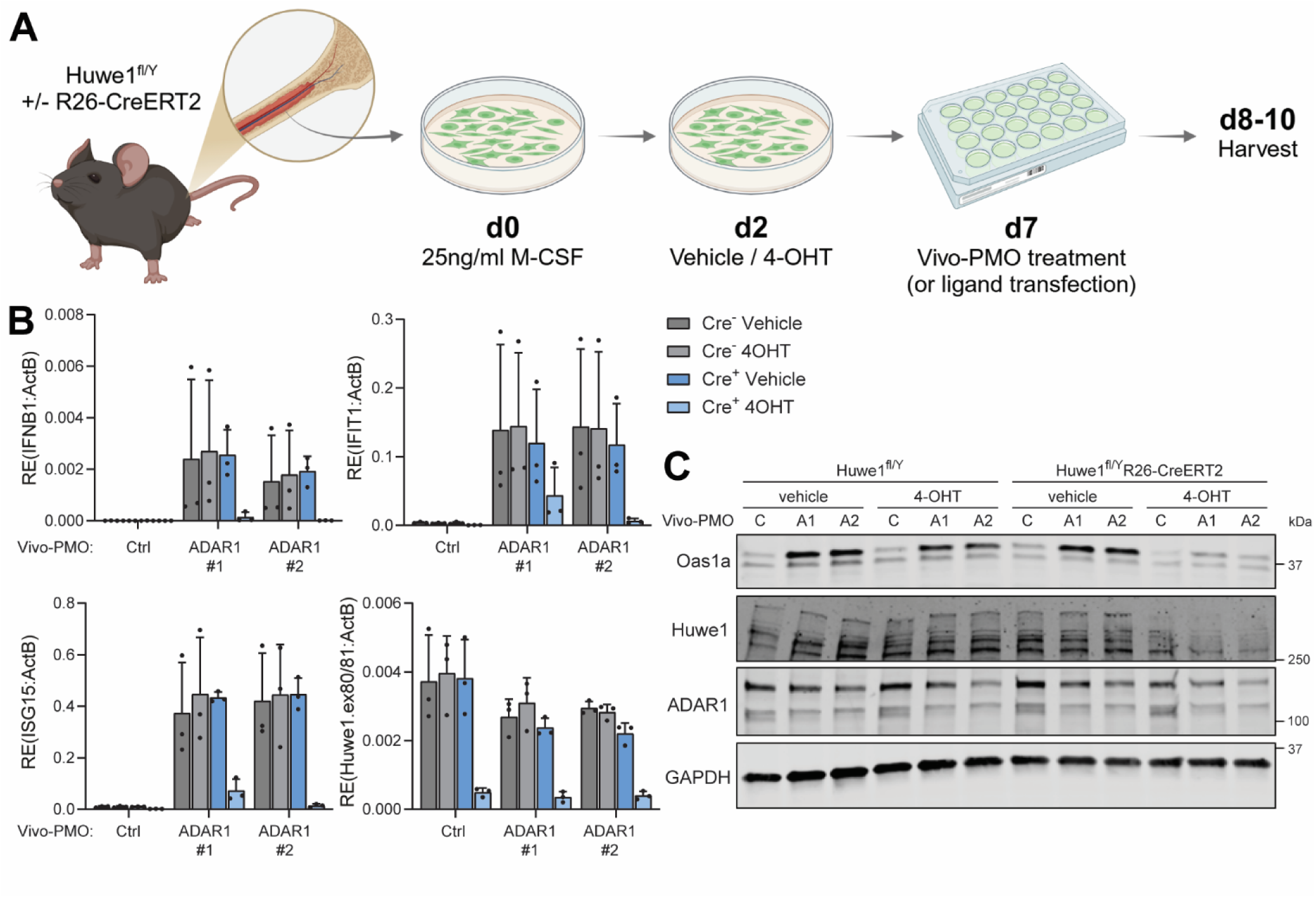
Conditional depletion of Huwe1 in primary BMDMs suppresses type I IFN induction upon loss of ADAR1. **A)** Schematic representation of experimental workflow. Bone marrow was collected from Huwe1^fl/Y^ (Cre^-^) and Huwe1^fl/Y^R26-CreERT2 (Cre^+^) mice and differentiated for 7 days with M-CSF to generate bone-marrow derived macrophages (BMDMs). On day 2 of differentiation, tamoxifen (4-OHT) was added to the cultures to induce Cre recombinase expression and Huwe1 depletion. A DMSO control (vehicle) is included as negative control. Cells were reseeded on day 6 and transfected on day 7 with indicated reagents to induce RLR activation. **B-C)** Differentiated BMDMs, generated as described in (A), were transfected with cell-penetrating phosphoroamidate morpholino oligomers (Vivo-PMOs) targeting ADAR1 or a control Vivo-PMO. Cells were harvested 72 h post-transfection. **B)** RT-qPCR analysis was used to monitor the type I IFN response and Cre-mediated recombination efficiency, using primers that anneal in the excised region of Huwe1 (exons 80-81). Transcripts were normalised to ACTB. **C)** Protein lysates were subjected to SDS-PAGE and immunoblotting using the indicated antibodies. Vivo-PMO C, Ctrl; A, ADAR1. Data are presented as means ± s.d. from n=3 experiments (B) or are representative data (n=3 for C).

### Identification of putative Huwe1 substrates upon RLR activation uncovers key components of the RLR pathway

To identify relevant Huwe1 substrates, we performed a proteome-wide analysis in Huwe1-sufficient and -deficient cells upon RLR stimulation. We made use of HEK293-L cells that were stably transduced with a lentivirus encoding a doxycycline (dox)-inducible shRNA targeting Huwe1 (i-shHuwe1), rather than the HEK293 Huwe1 KO cells, to avoid non-specific variations in the proteomic analysis inherently associated with single cell cloning. We validated that Huwe1 mRNA and protein levels were reduced in dox-treated i-shHuwe1 cells and not in i-shGFP control cells (Fig. S5A-D). Consistently, shRNA-mediated reduction of Huwe1 led to a reduced type I IFN response upon ADAR1 depletion or transfection of viral dsRNA (mimetics) (Fig. S5A-D), in line with above results (Fig. 1 and 3).

To determine the Huwe1-dependent ubiquitinome, we performed diGly remnant profiling^66^. This method relies on the immunoaffinity purification and MS-based detection of peptides with an unique Lys-ε-Gly-Gly motif (hereafter referred to as diGly peptides), generated by tryptic digestion of ubiquitinated proteins (Fig. 5A). Analyses were conducted in biological triplicates and a fraction of each sample was used for RT-qPCR analysis to confirm that the HMW poly(I:C)-induced type I IFN response was inhibited in dox-treated i-shHuwe1 cells, compared to untreated i-shHuwe1 cells (Fig. S5E). Total proteome analysis was included to investigate Huwe1-dependent changes in protein levels. A total of 7904 proteins and 4457 unique diGly peptides were identified. Principal component analysis (PCA) revealed robust clustering of the replicates from different treatments in the total proteome analysis, while replicate clusters were not as defined in the ubiquitinome analysis (Fig. S6A). Total proteome analysis revealed that several proteins known to be targeted by Huwe1 for proteasomal degradation (CHEK1, MAFB, MCL-1, and BRCA1)^50, 51, 67, 68^ were upregulated in Huwe1-depleted cells (Fig. S6B). Gene ontology (GO) enrichment analysis on proteins that were significantly up- or downregulated in Huwe1-depleted cells did not reveal an overrepresentation of proteins related to innate immune responses (Fig. S6B-C), suggesting that total protein levels of the RLR sensing machinery were largely unaffected upon loss of Huwe1. Virtually all proteins that were significantly upregulated by transfection of HMW poly(I:C) in control cells were known ISGs and their induction was largely reduced in poly(I:C)-treated Huwe1-depleted cells (Fig. S6D-E), confirming that Huwe1 is required for ISG upregulation upon RLR activation.

**Figure 5:**
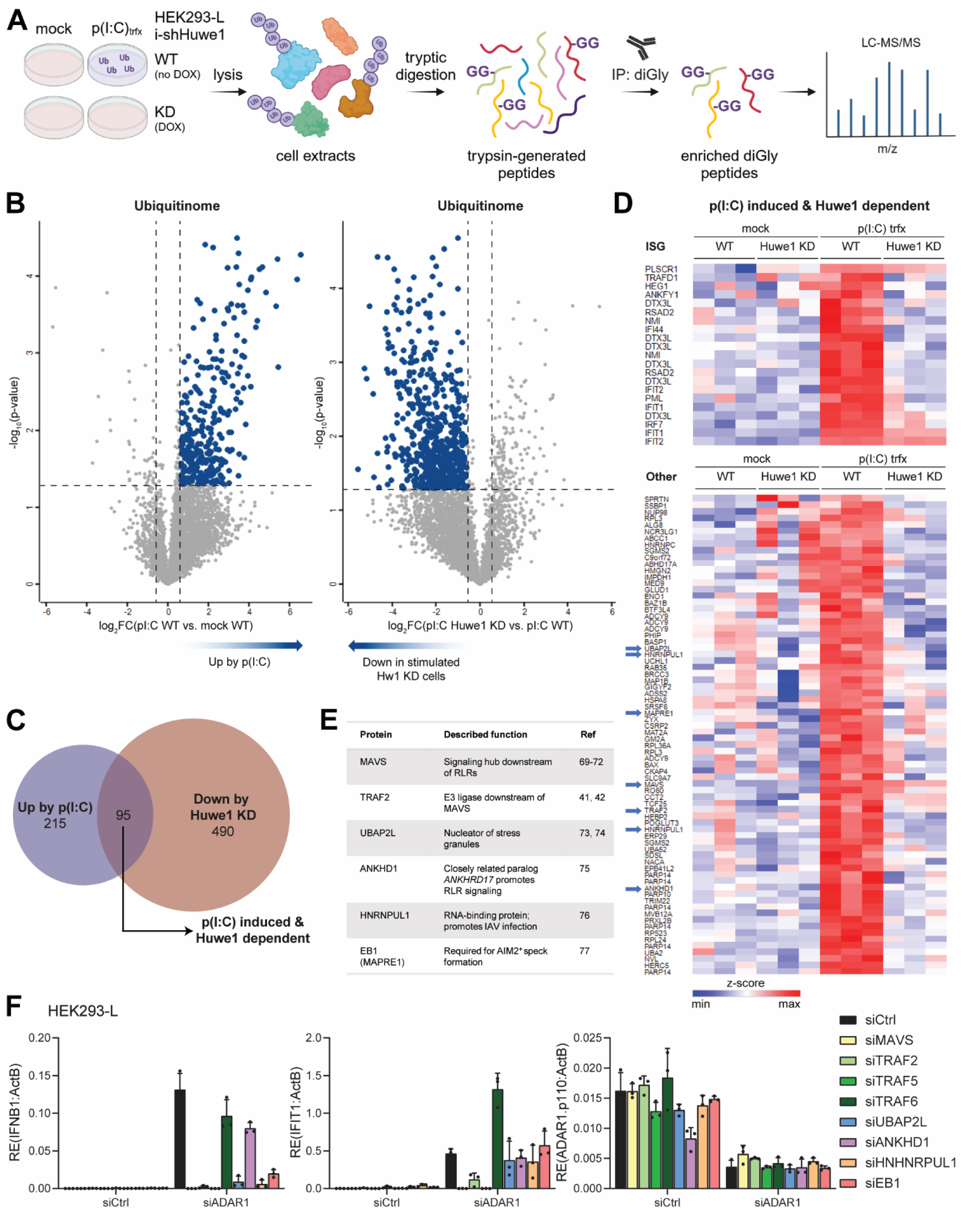
Ubiquitin diGly remnant profiling uncovers multiple potential Huwe1 substrates in the RLR pathway. **A)** Workflow for diGly remnant profiling. HEK293 cells transduced with doxycycline (DOX)-inducible shRNAs targeting Huwe1 (i-shHuwe1) were treated as indicated: -/+ DOX for 6 days followed by mock or high molecular weight poly(I:C) transfection (pI:C_trfx_) for 16 h. After tryptic digestion of cell lysates, Lys-ε-Gly-Gly containing (diGly) peptides were affinity purified and subjected to analysis by mass-spectrometry to acquire the ubiquitinome. **B)** Volcano plots comparing changes in the ubiquitinome due to poly(I:C) transfection of wild type (WT) cells (*left*), or due to Huwe1 KD under conditions of poly(I:C)_trfx_ (*right*), in n=3 independent experiments. Dashed lines indicate significance cutoff at a p-value of 0.05 (-log_10_ = 1.3) in y-axis and a fold change (FC) of 1.5 (log_2_FC = 0.6) in x-axis. **C)** Proportional Venn diagram of diGly peptides, highlighted in blue in (B), that are significantly upregulated by poly(I:C) transfection in WT cells or downregulated due to Huwe1 KD in poly(I:C) transfected cells. Their overlap reveals diGly peptides that are poly(I:C) induced and Huwe1 dependent. **D)** Heat maps of poly(I:C)-induced and Huwe1 dependent diGly peptides (95 overlapping peptides in C), divided into peptides that originate from ISGs and other proteins. Intensity values were normalized by Z score. Peptides belonging to putative Huwe1 substrates that were selected for follow-up analysis are indicated with blue arrows. **E)** Summary of putative Huwe1 substrates that were selected for follow-up analysis. **F)** HEK293 cells that stably express FLAG-LGP2 (HEK293-L) were transfected with the indicated siRNAs and harvested 72 h post-transfection. RT-qPCR analysis was used to monitor the type I IFN response and ADAR1 knockdown efficiency. Transcripts were normalised to ACTB. Data are presented as means ± s.d. from n=3 experiments.

The ubiquitinome of the different treatment groups were cross-compared to identify ubiquitination events that are induced upon HMW poly(I:C) transfection in a Huwe1-dependent manner. This revealed 95 diGly peptides that were both increased upon HMW poly(I:C) transfection and reduced in poly(I:C)-treated Huwe1-depleted cells (Fig. 5B-C). A number of these peptides were derived from ISGs, which were likely detected due to their induction upon RLR activation in control but not Huwe1-depleted cells (Fig. 5B). The majority of peptides were, however, not derived from ISGs (Fig. 5B).

Six putative Huwe1 substrates were selected for follow-up analysis (Fig. 5E). Total protein levels of these targets were, except for HNRNPUL1, largely unaffected by the loss of Huwe1 (Fig. S6F). Of particular interest were MAVS and TRAF2, as these proteins are central components of the RLR pathway. Functional similarities and redundancies between TRAF2 and other TRAF family members, in particular TRAF5, in promoting RLR-induced IFN responses have been reported^41, 42^. Hence, TRAF5 was also included in follow-up analysis. TRAF6 was also included, as Huwe1 has previously been reported to regulate TRAF6-dependent IL-1β signalling^57^. The other selected targets (Fig. 5E) are not integral to the RLR pathway and may have an indirect impact on RLR signalling. We systematically assessed for each of these targets whether its expression is required for type I IFN induction and RLR signalling following loss of ADAR1. As expected, loss of the central RLR signalling hub MAVS abrogated type I IFN production (Fig. 5D; Fig. S6G). Loss of TRAF2 or TRAF5, but not TRAF6, also hampered the type I IFN response to unedited self RNA. IFN-β expression was also somewhat reduced in cells depleted of UBAP2L, HNRNPUL, or MAPRE1, although IFIT1 expression was less affected. Collectively, we find that RLR activation promotes Huwe1-dependent ubiquitination of multiple proteins that aid in type I IFN production and that loss of MAVS, TRAF2, and TRAF5 had the most notable impact on type I IFN induction following loss of ADAR1.

### Huwe1 promotes TRAF5-mediated type I IFN responses in concert with UBR5

To substantiate the functional interplay between Huwe1 and TRAFs or MAVS, we performed co-immunoprecipitation experiments. Ectopically expressed Huwe1 interacted with TRAF2, TRAF5, TRAF6 and MAVS (Fig. 6A), revealing that Huwe1 physically associates with these central components of the RLR pathway. Curiously, expression of TRAF2, TRAF5, or TRAF6 enhanced protein expression of ectopic, but not endogenous, Huwe1 (Fig. 5A-B). We next analysed how Huwe1 impacts on the type I IFN response induced by overexpression of the different TRAFs or MAVS. We observed that induction of the type I IFN response upon TRAF5 overexpression and autoactivation was largely reduced in Huwe1-depleted cells, whereas Huwe1 did not impact on MAVS-induced IFN induction (Fig. 6B). Expression of TRAF2, TRAF3, and TRAF6 did not induce a spontaneous type I IFN response, regardless of Huwe1 expression. Under conditions where ectopic expression of TRAF5 induces ISG expression, endogenous Huwe1 interacted with TRAF5 (Fig. 6C). These data indicate that Huwe1 facilitates the type I IFN response either directly or indirectly via TRAF5 activity.

**Figure 6:**
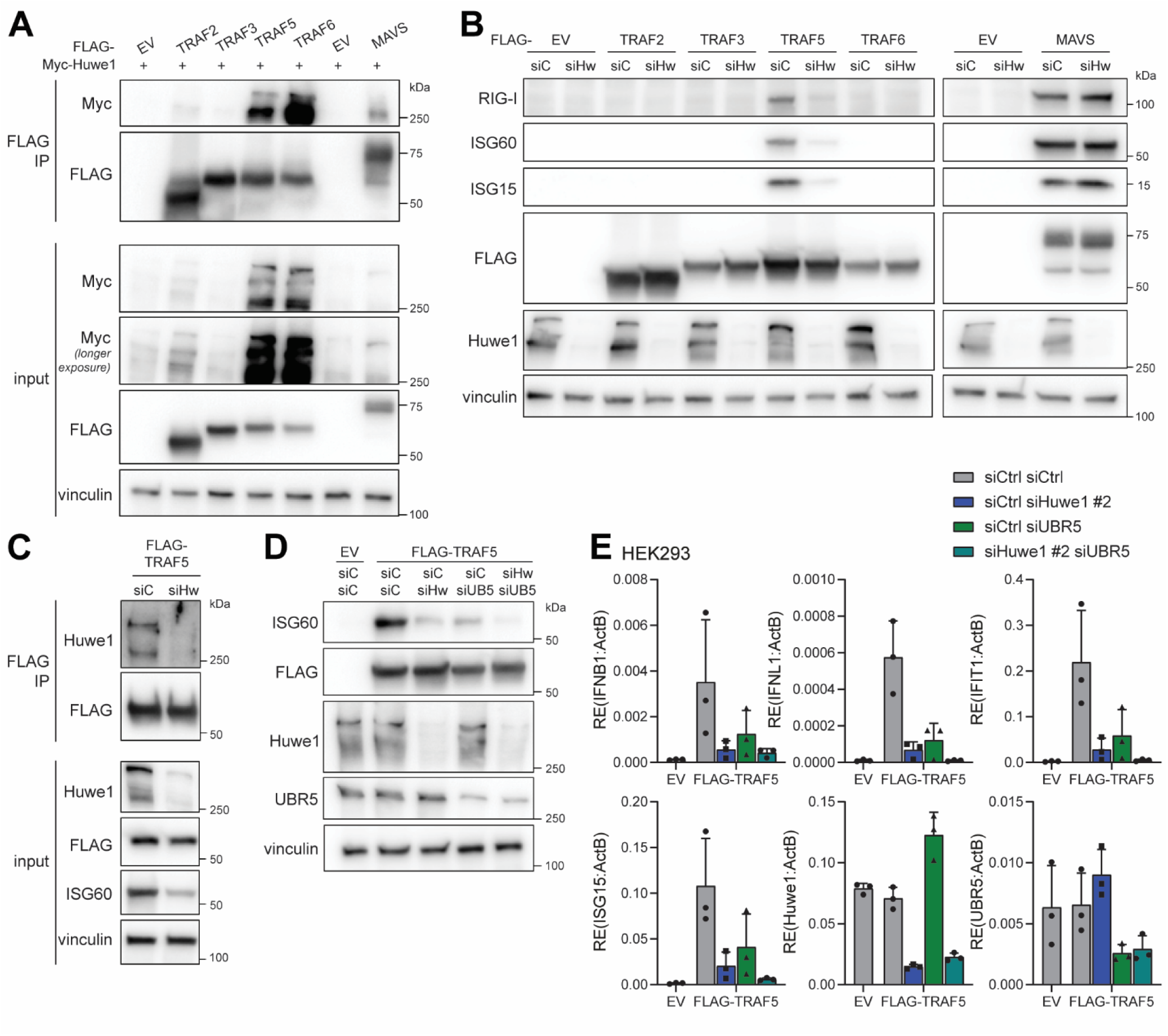
Huwe1 is required for type I IFN induction via TRAF5 and works in concert with UBR5. **A)** Co-immunoprecipitation analysis to assess the interaction of ectopically expressed Myc-Huwe1 with the indicated FLAG-tagged proteins. Protein lysates of transfected HEK293 cells were subjected to FLAG immunoprecipitation (IP). IP (top) and input (bottom) fractions were subjected to SDS-PAGE and immunoblotting using the indicated antibodies. EV, empty vector. **B)** HEK293 cells were transfected with the indicated siRNAs and 24 h later with an empty vector (EV) or vector encoding the indicated FLAG-tagged protein. Samples were harvested 72 h post siRNA-transfection and subjected to SDS-PAGE and immunoblotting using the indicated antibodies. siC, siCtrl; siHw, siHuwe1 #2. **C)** Co-immunoprecipitation analysis to assess the interaction of endogenous Huwe1 with ectopically expressed FLAG-TRAF5. Protein lysates of transfected HEK293 cells were subjected to FLAG immunoprecipitation (IP). IP (top) and input (bottom) fractions were subjected to SDS-PAGE and immunoblotting using the indicated antibodies. siC, siCtrl; siHw, siHuwe1 #2. **D-E)** HEK293 cells were transfected with the indicated siRNAs and 24 h later with an empty vector (EV) or a vector encoding the indicated FLAG-tagged protein. Samples were harvested 72 h post siRNA-transfection. **D)** Protein lysates were subjected to SDS-PAGE and immunoblotting using the indicated antibodies. siC, siCtrl; siHw, siHuwe1 #2, siUB5, siUBR5. **E)** RT-qPCR analysis was used to monitor the type I and type III IFN response and knockdown efficiencies of Huwe1 and UBR5. Transcripts were normalised to ACTB. Data are presented as means ± s.d. from n=3 experiments (E) or are representative data (n=3 for A, C, D; n=2 for B).

Existing evidence suggests that Huwe1 may collaborate with other giant HECT-domain containing ubiquitin E3 ligases, such as UBR5^78–80^, through overlapping substrate specificities. We compared how siRNA-mediated knockdown of Huwe1 and UBR5, either individually or simultaneously, affects the TRAF5-induced type I IFN response. Depletion of Huwe1 or UBR5 alone partially reduced expression of type I and III IFNs as well as ISGs, while the combined depletion of both Huwe1 and UBR5 completely abrogates the TRAF5-induced type I IFN response (Fig. 6D-E). Altogether, these data suggest that Huwe1, in conjunction with UBR5, facilitates type I IFN induction at least in part by regulating TRAF5 activity.

## Discussion

In summary, we have demonstrated an important role for the ubiquitin E3 ligase Huwe1 in the induction of a type I and type III IFN response following the activation of RLRs. We find that Huwe1 plays a role both in context of viral and endogenous RNA sensing in a range of cell types. Through diGly remnant profiling, we have determined how the loss of Huwe1 alters the ubiquitinome during RLR activation. We have identified a number of potential substrates that interact with Huwe1, including TRAFs and MAVS. Strikingly, we find that the activity of TRAF5 is Huwe1-dependent, suggesting that TRAF5 is a prime candidate via which Huwe1 may exert its regulatory role in the RLR pathway.

The impact of Huwe1 on type I IFN induction via RLRs differs in magnitude depending on the context. We find that, irrespective of the type of depletion method used, loss of Huwe1 nearly completely abrogates type I IFN induction in response to the accumulation of endogenous immunostimulatory RNA in ADAR1-deficient cells. In contrast, upon activation of RLRs by viral RNA mimetics or RNA virus infection, Huwe1 depletion causes a more modest reduction in type I IFN induction. The importance of Huwe1 in type I IFN induction may be dictated by cell type, where cells of epithelial and mesenchymal origin are more dependent on the presence of Huwe1 than macrophages. In addition, the detection of viral RNA mimetics was modestly dependent on Huwe1 in human monocyte-derived macrophages and independent of Huwe1 in murine BMDMs, pointing to species-specific differences. The extent by which Huwe1 impacts on RLR-mediated type I IFN production may depend on cell-type specific expression levels and redundancy of TRAF proteins, or a different requirement for TRAF proteins in sensing of viral versus endogenous RNA ligands. On a more technical note, we noticed that Huwe1 is visible as multiple species of distinct size by immunoblot; while siRNA- or shRNA-mediated knockdown of Huwe1 in human cells led to a complete loss of all species, tamoxifen-treated BMDMs derived from Huwe1^fl/Y^R26-CreERT2 mice have only lost the upper species.

While this study focused on type I IFN induction following RLR activation, our data suggest that Huwe1 may also impact on cGAS/STING signaling upon activation by cytosolic dsDNA. We did not explore whether this effect is mediated via TRAF proteins or via an independent mechanism. The role of TRAF proteins (including TRAF5) in the cGAS/STING pathway remains poorly defined. TRAF3 may be activated by STING upon infection with human enterovirus A71 and is a target for cleavage by the viral protease 2A^pro 81^. In addition, TRAF6 has been implicated in cGAS/STING signaling upon infection with herpes simplex virus and the C-terminus of STING contains putative TRAF2-binding motifs^82, 83^. Additional studies in this underexplored area may provide insight into the precise role of different TRAF proteins in the cGAS/STING pathway, and whether these are regulated in a Huwe1-dependent manner. On a similar note, it would be interesting to explore whether Huwe1 also impact on TRAF5 activity in other TRAF5-dependent pathways, such as IL-17-mediated signaling^84, 85^.

It remains unclear whether Huwe1 directly targets TRAF5 for ubiquitination, or whether it regulates TRAF5 indirectly, for example via other TRAF proteins. Since TRAF proteins, which are ubiquitin E3 ligases themselves, may undergo autoubiquitination^86–89^, it is technically challenging to determine how the absence or presence of Huwe1 alters their ubiquitination status. In addition, Huwe1 may exert its role in type I IFN induction not exclusively via TRAF5, but in a multimodal manner and via multiple substrates and pathways. Interestingly, Huwe1 has recently been implicated in the clearance of protein aggregates, including G3BP1-positive stress granules ^90^. The precise role of RNA granules, including stress granules, in nucleic acid sensing remains a matter of ongoing debate, with some seemingly conflicting reports^91–93^. Amongst our potential Huwe1 substrates we identified several proteins that participate in stress granule formation. An attractive hypothesis, and an ongoing line of investigation in the lab, is that Huwe1 controls the emergence or persistence of stress granules via the regulation of stress granule nucleators, and thereby impacts on nucleic acid sensing.

Our findings support a key role for Huwe1 in promoting type I IFN induction following RLR activation, consistent with other reports that implicate Huwe1 as a positive regulator of other aspects of innate immune activation and inflammatory signalling. RNA sequencing revealed that loss of Huwe1 leads to a repression of genes involved in innate immunity and inflammatory signalling^68^. In addition, Huwe1 depletion reduces NF-κB activation via TRAF6^57^. Consistently, loss of Huwe1 led to reduced expression of IL-6 and TNFα transcripts upon LPS stimulation of murine macrophages^94^. Finally, IL-1β secretion was reduced in Huwe1-deficient BMDMs and mice in response to inflammasome-activating ligands and upon infection with intracellular bacteria^58^. Together with our own observations, these studies contribute to the emerging view that Huwe1 plays an important role in enhancing inflammation. Interestingly, Huwe1 was a target of viral antagonism by two distinct respiratory viruses (SARS-CoV-2 and Influenza A virus) albeit via distinct mechanisms^95, 96^. Altered Huwe1 protein expression correlated with a reduced antiviral response and increased viral replication. The notion that Huwe1 is a potential immune evasion target underscores its importance in antiviral innate immune signalling.

Huwe1 belongs to a class of giant HECT-domain containing E3 ligases that do not only extend polyubiquitin chains but also generate branched ubiquitin polymers (which is sometimes referred to as ‘E4 ligase activity’)^97^. Huwe1 catalyzes K48-linked branches onto K63-linked chains formed by TRAF6 to prolong NF-κB activation^57^. The assembly of branched ubiquitin chains generally involves the cooperation between multiple E3 ligases with different linkage specificities. In this context, Huwe1, UBR4, and UBR5 may collaborate to catalyze such branched linkages^98^. These E3 ligases were associated with a number of shared substrates, including TXNIP, Apobec3 proteins, TRIM52, and Orf9b (from SARS-CoV-2)^78–80, 99^. In addition, both UBR5 and Huwe1 interact with K11/K48-branched chains^100^. We find that the simultaneous depletion of both Huwe1 and UBR5 limits TRAF5-induced type I IFN expression more strongly than knockdown of each individual component. It will be worthwhile to investigate whether the regulation of TRAF5 or other potential Huwe1 substrates involves branched ubiquitin chain formation.

In conclusion, we have uncovered an important role for the ubiquitin E3 ligase Huwe1 in promoting RLR-mediated type I and type III IFN induction. These findings highlight the importance of ubiquitin-dependent regulation of cytosolic RNA sensing and the downstream type I and type III IFN response.

---

**Supplemental Figure 1.**
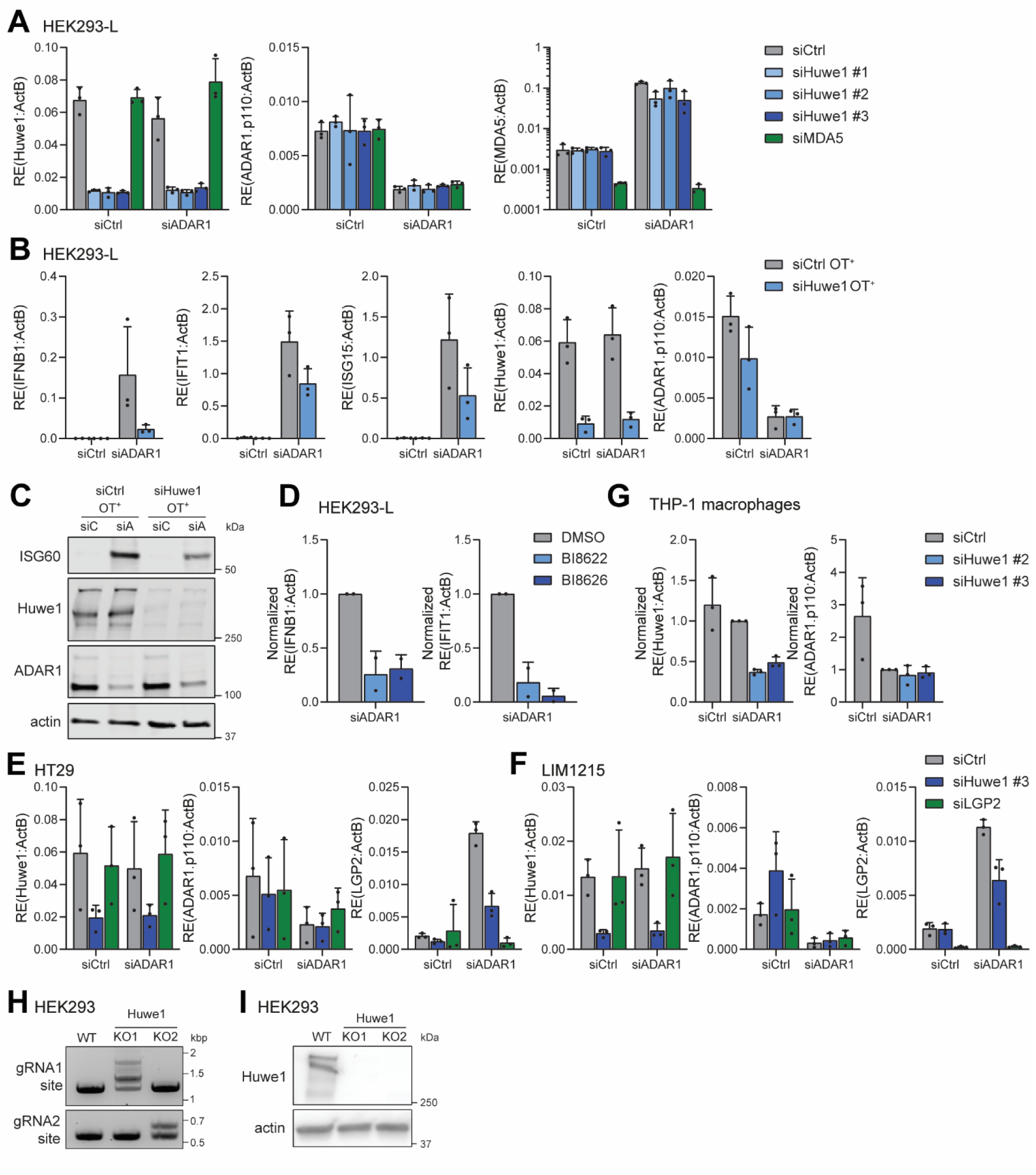
, belonging to Figure 1 **A)** RT-qPCR analysis was used to monitor the efficiency of siRNA-mediated knockdown of Huwe1, ADAR1, and MDA5 in samples of HEK293 cells that stably express FLAG-LGP2 (HEK293-L) from Fig. 1A. **B-C)** HEK293-L cells were transfected with combinations of a control, ADAR1- or Huwe1-targeting pool of ON-TARGETplus (OT+) siRNAs. Samples were harvested 72 h post-transfection. **B)** RT-qPCR analysis was used to monitor the type I IFN response and the efficiency of Huwe1 and ADAR1 knockdown. Transcripts were normalised to ACTB. **C)** Protein lysates were subjected to SDS-PAGE and immunoblotting using the indicated antibodies. siC, siCtrl; siA, siADAR1. **D)** HEK293-L cell were transfected with siADAR1. At 16h and 48h post siRNA-transfection medium was replaced with fresh medium supplemented with 20 µM of the Huwe1-inhibitor BI8622, 25 µM of the inhibitor BI8626, or a vehicle control (DMSO). RT-qPCR analysis was used to monitor the type I IFN response. Transcripts were normalised to ACTB and are displayed relative to the vehicle control. **E-F)** RT-qPCR analysis was used to monitor the efficiency of siRNA-mediated knockdown of Huwe1, ADAR1, and LGP2 in samples of HT29 and LIM1215 cells from Fig. 1D and E, respectively. **G)** RT-qPCR analysis was used to monitor the efficiency of siRNA-mediated knockdown of Huwe1 and ADAR1 in samples of PMA-differentiated THP-1 macrophages from Fig. 1F. **H-I)** HEK293 cells were genetically engineered to knockout Huwe1 via CRISPR/Cas9. Two Huwe1 knockout (KO) clones were expanded, that were generated using individual gRNAs that target distinct regions in the Huwe1 gene (KO1 = gRNA1; KO2 = gRNA2). **H)** Primers flanking either gRNA target site in the Huwe1 gene were used to confirm targeted indel generation in the Huwe1 gene by PCR analysis. **I)** Immunoblotting confirmed the loss of Huwe1 protein in the Huwe1 KO HEK293 clones. Data are presented as means ± s.d. from n=3 (A, B, E-G) or n=2 (D) experiments or are representative data (n=2 for C, H; n=3 for I).

**Supplemental Figure 2.**
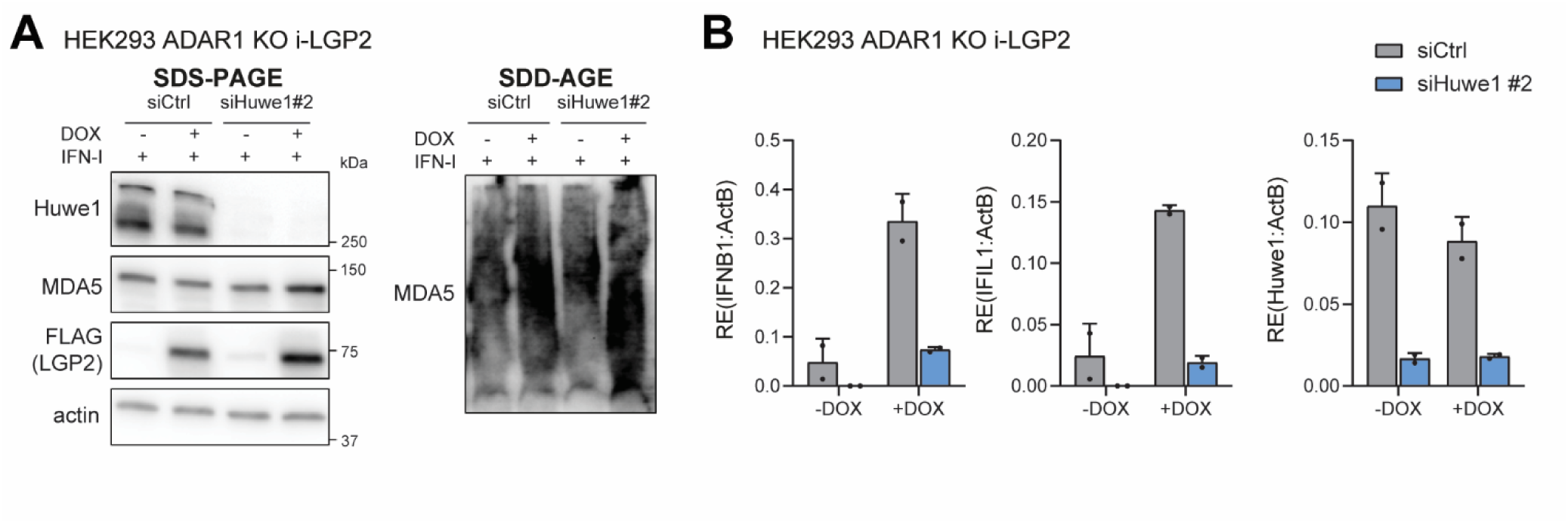
, belonging to Figure 2 **A-B)** ADAR1-knockout (KO) HEK293 cells transduced with a lentiviral-based inducible system to express FLAG-LGP2 in a doxycycline (DOX)-regulated manner (i-LGP2) were transfected with the indicated siRNAs and treated +/- DOX for 72 h. **A)** During the last 24 h, recombinant type I IFN (1000 U/ml; IFN-I) was added to upregulate endogenous MDA5 protein expression. Protein lysates were subjected to SDD-AGE and SDS-PAGE, followed by immunoblotting using the indicated antibodies, to determine MDA5 oligomerization and total protein levels, respectively. Representative immunoblots are shown (n=2). **B)** RT-qPCR analysis was used to monitor the type I and type III IFN response and Huwe1 knockdown efficiency. Transcripts were normalised to ACTB. Data are presented as means ± s.d. from n=2 experiments.

**Supplemental Figure 3.**
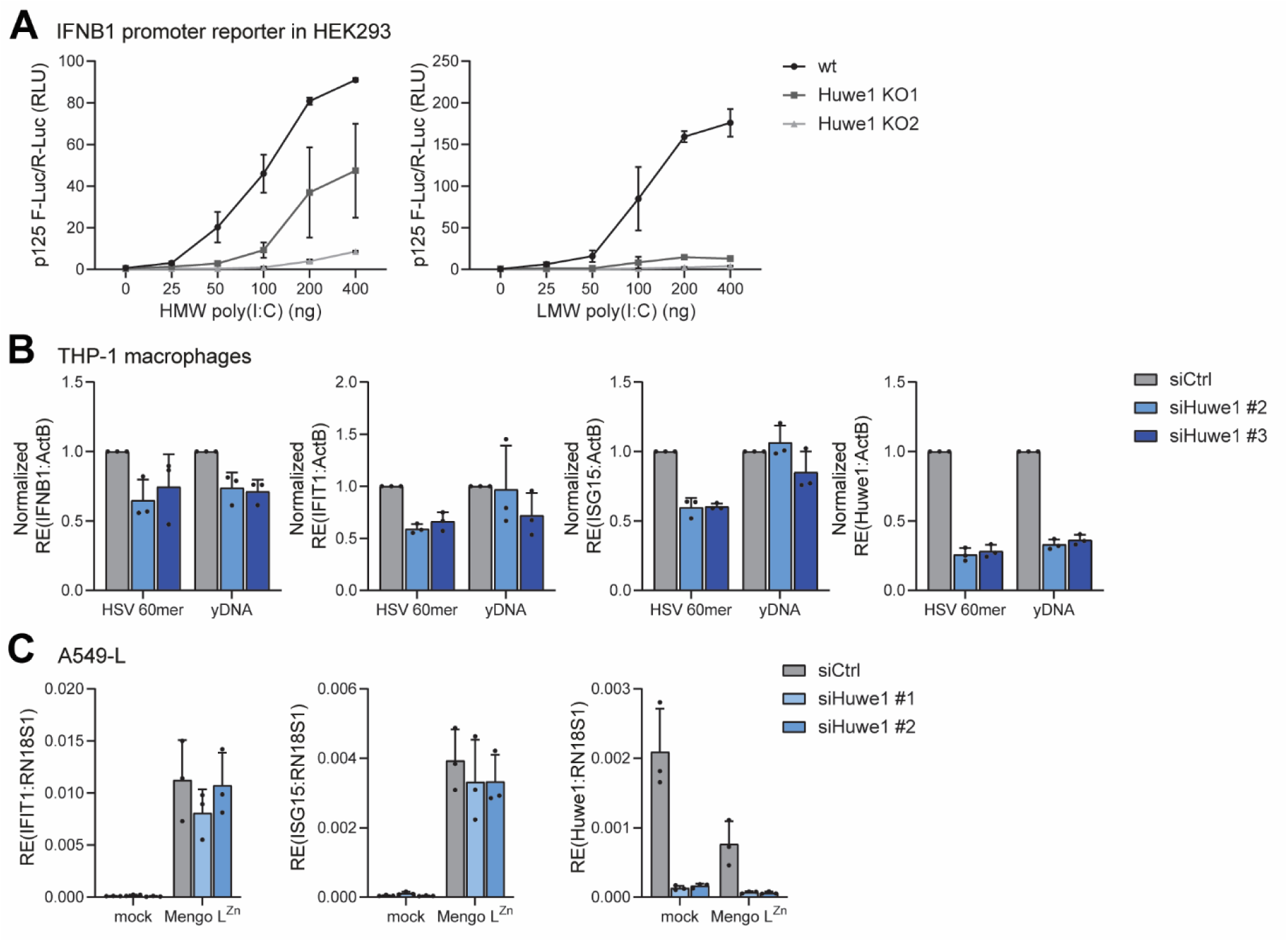
, belonging to Figure 3 **A)** Wild type (WT) and Huwe1 knockout (KO) HEK293 cells were transfected with p125-luc (IFN-β promoter reporter) plasmid and Renilla luciferase vector (internal control). After 24 h, cells were transfected with the indicated dose of high molecular weight (HMW) or low molecular weight (LMW) poly(I:C), and after overnight incubation, luciferase activities were determined. **B)** PMA-differentiated THP-1 macrophages were transfected with the indicated siRNAs. After 64 h, cells were transfected for an additional 6 h with 4 µg of the dsDNA ligands HSV 60mer or G3-YSD (yDNA). RT-qPCR analysis was used to monitor the type I IFN response and Huwe1 knockdown efficiency. Transcripts were normalised to ACTB and are displayed relative to siCtrl. **C)** RT-qPCR analysis was used to monitor ISG expression and the efficiency of siRNA-mediated knockdown of Huwe1 in samples of A549 cells that stably express FLAG-LGP2 (A549-L) from Fig. 3C. Data are presented as means ± s.d. from n=2 (A) or n=3 (B, C) experiments.

**Supplemental Figure 4.**
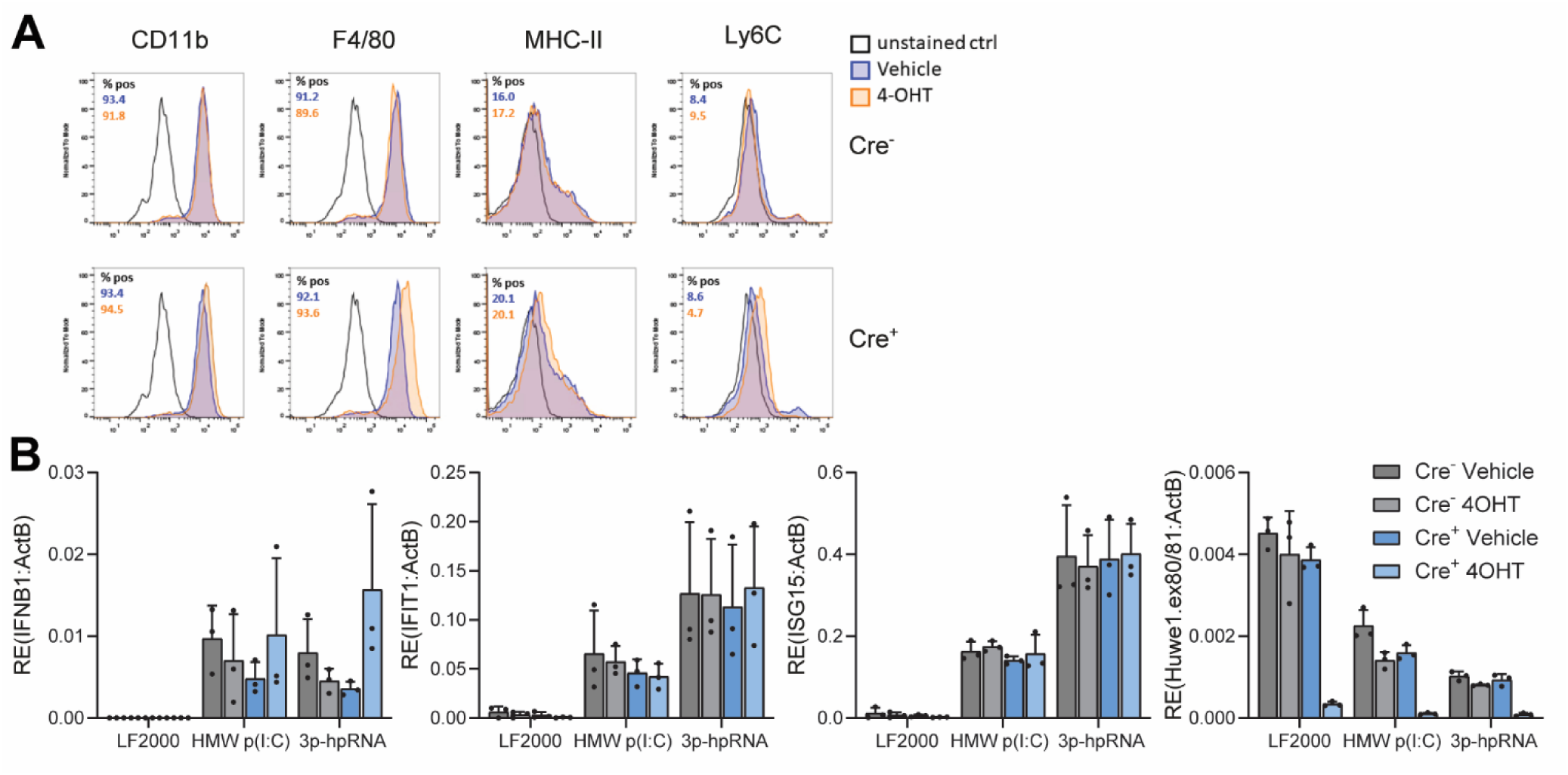
, belonging to Figure 4 **A)** BMDMs were generated from bone marrow of Huwe1^fl/Y^ and Huwe1^fl/Y^R26-CreERT2 mice and treated with a vehicle control or 4-OHT as depicted in Fig. 4A. The surface levels of the indicated differentiation markers were determined using flow cytometry (n=1). **B)** Differentiated BMDMs, generated as described in (Fig. 4A), were transfected with transfection reagent only (LF2000), 0.5 µg high molecular weight (HMW) poly(I:C), or 0.3 µg of a small triphosphorylated hairpin RNA (3p-hpRNA). Samples were harvested 16 h post-transfection and RT-qPCR analysis was used to monitor the type I IFN response and Cre-mediated recombination efficiency, using primers that anneal in the excised region of Huwe1 (exons 80-81). Transcripts were normalised to ACTB. Data are presented as means ± s.d. from n=3 experiments.

**Supplemental Figure 5.**
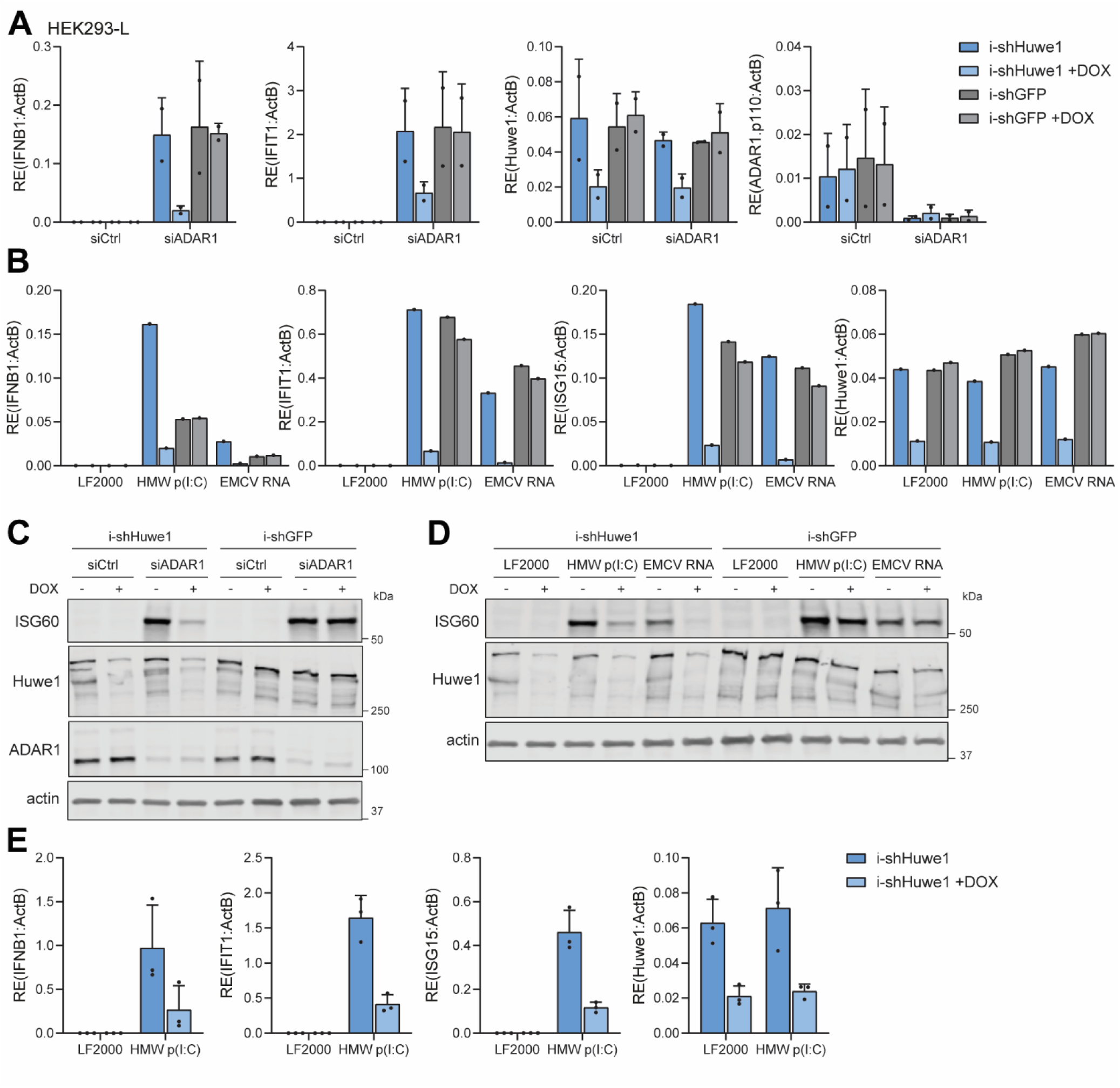
, belonging to Figure 5 **A-D)** HEK293 cells that stably express FLAG-LGP2 (HEK293-L) and are transduced with doxycycline (DOX)-inducible shRNAs targeting Huwe1 (i-shHuwe1) or GFP (i-shGFP; negative control) were treated +/- DOX for 7 days. Cells were transfected with the indicated siRNAs for the last 72 h (A and C) or for the last 16 h with indicated ligands that activate MDA5 (B and D). **A-B)** RT-qPCR analysis was used to monitor the type I IFN response and the efficiency of Huwe1 and ADAR1 knockdown. Transcripts were normalised to ACTB. **C-D)** Protein lysates were subjected to SDS-PAGE and immunoblotting using the indicated antibodies. **E)** RT-qPCR analysis was used to monitor the type I IFN response and the efficiency of Huwe1 and ADAR1 knockdown in samples used for diGly profiling (Fig. 5). Transcripts were normalised to ACTB. Data are presented as means ± s.d. (n=2 for A, n=1 for B, n=3 for E) or are representative data (n=2 for C; n=1 for D).

**Supplemental Figure 6.**
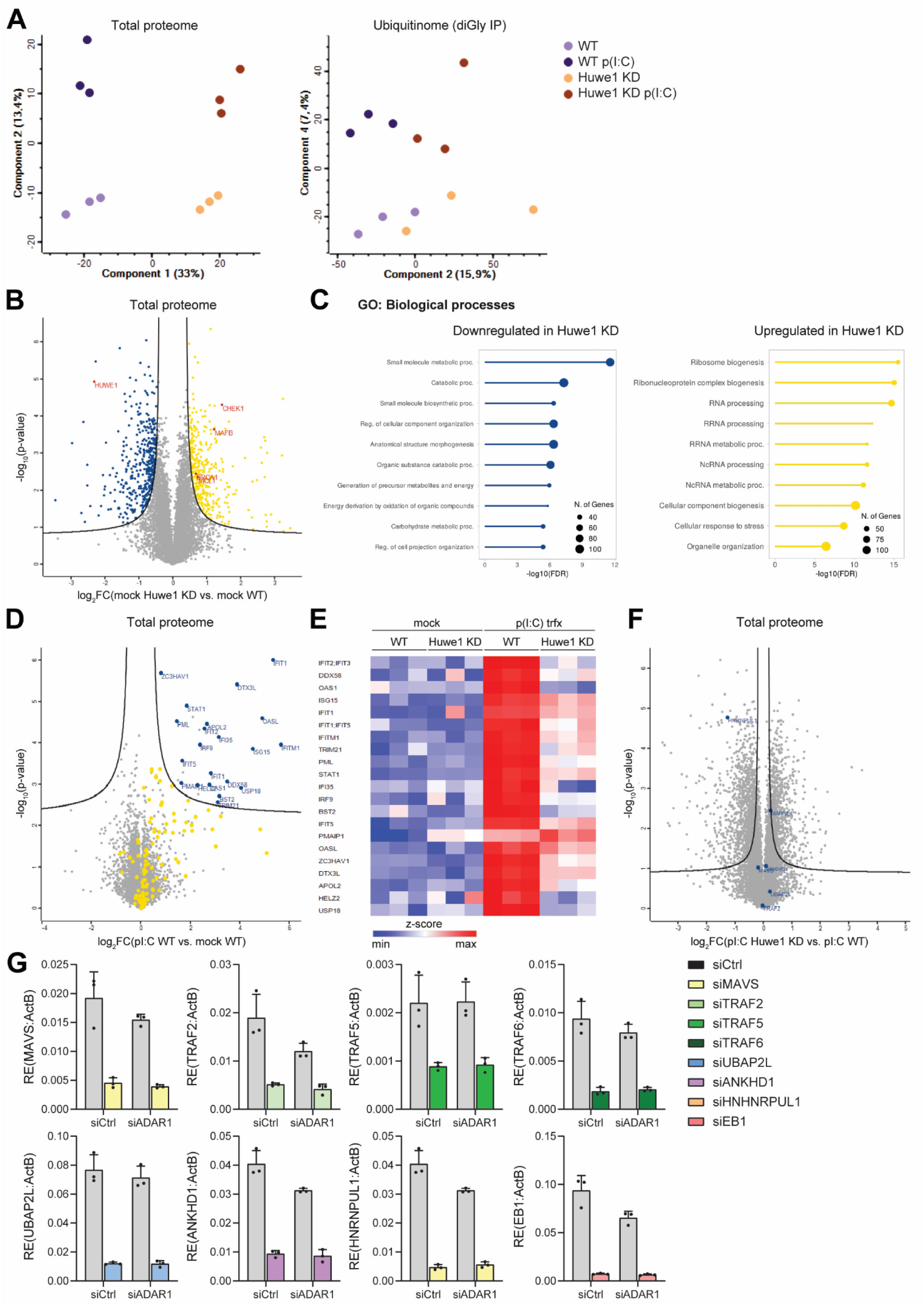
, belonging to Figure 5 **A)** Principal components analysis (PCA) of the total proteome and ubiquitinome (diGly profiling) data-independent acquisition (DIA) data. **B)** Volcano plot comparing changes in the total proteome due to Huwe1 knockdown (KD), in n=3 independent experiments (statistical parameters: FDR = 0.05; S0 = 0.25). Significantly downregulated (blue) and upregulated (yellow) proteins are indicated. The loss of Huwe1 and upregulation of known targets of Huwe1-induced proteasomal degradation are highlighted in red. **C)** GO enrichment analysis of proteins that are significantly downregulated (blue) or upregulated (yellow) upon Huwe1 KD (as defined in B). Graphs show the most significantly overrepresented biological processes. **D)** Volcano plot comparing changes in the total proteome due to poly(I:C) transfection in WT cells, in n=3 independent experiments (statistical parameters: FDR = 0.05; S0 = 0.1). Known ISGs^101, 102^ were highlighted in yellow and significantly upregulated ISGs are shown in blue. **E)** Heat map ISGs that were significantly upregulated by poly(I:C) transfection in WT cells (as defined in D), reveals reduced ISG upregulation in poly(I:C)-treated Huwe1 KD cells. Intensity values were normalized by Z score. **F)** Volcano plot comparing changes in the total proteome between Huwe1 KD and WT cells under conditions of poly(I:C) transfection, in n=3 independent experiments (statistical parameters: FDR = 0.05; S0 = 0.1). Putative Huwe1 substrates selected for follow-up analysis (Fig. 5E) are highlighted in blue. **G)** RT-qPCR analysis was used to monitor the efficiency of siRNA-mediated knockdown in samples of HEK293 cells that stably express FLAG-LGP2 (HEK293-L) from Fig. 5F. Data are presented as means ± s.d. from n=3 experiments.

